# Genetic and epigenetic diversity of *Salmonella enterica* isolates from Kazakhstan from clinical and veterinary sources

**DOI:** 10.64898/2026.06.01.729238

**Authors:** Duman T. Yessimseit, Altyn K. Rysbekova, Zauresh B. Zhumadilova, Beck Z. Abdeliyev, Nur B. Tukhanova, Altynai K. Kassenova, Aigul K. Abdrakhmanova, Akmaral Mereke, Sanzhar D. Agzam, Ainur S. Nurpeisova, Raikhan Nissanova, Ayaulym M. Maksatova, Oleg N. Reva, Aigul A. Abdirassilova

**Affiliations:** M. Aikimbayev’s National Scientific Center for Especially Dangerous Infections, 14 Zhakhanger St., Almaty A35P0K3, Kazakhstan; Al-Farabi Kazakh National University, 71 al-Farabi Ave., Almaty A15E3B4, Kazakhstan; I. Zhekenova’s City Clinical Infectious Disease Hospital, Almaty A36G2X0, Kazakhstan; Kazakh Scientific Research Veterinary Institute, 223 Rayymbek Ave., Almaty A20C2E4, Kazakhstan; Centre for Bioinformatics and Computational Biology, Department of Biochemistry, Genetics and Microbiology, University of Pretoria, Pretoria 0002, South Africa

**Keywords:** *Salmonella enetrica*, genome, virulence, antibiotic resistance, plasmid, epigenetics

## Abstract

**Background:** *Salmonella enterica* is a major cause of foodborne and invasive infections worldwide. Increasing antimicrobial resistance and adaptation to diverse ecological niches require an improved understanding of the genetic and epigenetic diversity of circulating strains. This study investigated the genomic and epigenetic diversity of *S. enterica* isolates collected in Kazakhstan from clinical, animal, and environmental sources.

**Methods:** Whole-genome sequencing was performed using the Illumina sequencing platform. Several selected strains were additionally sequenced using PacBio SMRT technology for DNA methylation profiling. Genome assembly, plasmid reconstruction, MLST genotyping, and analyses of virulence genes, antimicrobial resistance determinants, and genome methylation associated with restriction-modification (RM) systems and orphan methyltransferases were performed using established bioinformatics tools.

**Results:** The ST11 genotype predominated among clinical isolates, but these strains formed distinct clusters differing in plasmid composition, virulence-associated genes, and resistance determinants. Most strains carried two large plasmids associated with environmental persistence and virulence, whereas the recent hospital isolate 19S, belonging to the ST11 group, carried two alternative plasmids enriched in virulence and antibiotic resistance genes. All genomes demonstrated conserved DAM-associated adenine methylation at *GATC* motifs, partial DCM-mediated cytosine methylation at *CCWGG* motifs, and widespread adenine methylation at *CAGAG* motifs linked to type III RM system. In contrast, the type I RM system present in the majority of sequenced strains was suppressed under laboratory growth conditions and remained active only in strain 19S, possibly due to mutations identified in the *hsdM* gene that may have released this methyltransferase from suppression. Novel epigenetic modification signals involving cytosine and guanine in replichore-biased tandem repeats were also identified.

**Conclusions:** *S. enterica* strains circulating in Kazakhstan exhibit substantial genomic and epigenetic diversity associated with different survival and transmission strategies. DNA methylation profiling provided additional insights beyond conventional MLST genotyping and identified strain 19S as a promising model for future studies of epigenetic regulation in bacterial virulence and adaptation mediated through genomic DNA methylation.

## Introduction

According to Worldwide Health Organization (WHO), foodborne infections caused by bacteria of the genus *Salmonella* remain a major public health concern worldwide because of their broad host range, high transmission potential, and increasing levels of antimicrobial resistance [1]. The expectation that climate warming may promote the widespread dissemination of antibiotic-resistant pathogens has recently been supported by studies using *Salmonella* as a model organism [2]. This finding highlights the need to establish surveillance measures for dynamic monitoring of changes in pathogen populations and the emergence of new unusual genotypes.

Members of this genus belong to the family Enterobacteriaceae and are currently classified into two species, *Salmonella bongori* and *Salmonella enterica* [3]. Among them, *S. enterica* is responsible for the overwhelming majority of infections in humans and warm-blooded animals. The species is highly heterogeneous and includes numerous lineages adapted to different ecological niches and transmission strategies [4].

Within *S. enterica*, the subspecies *enterica* contains most clinically important isolates and is further subdivided into a large number of serovars based on combinations of somatic and flagellar antigens together with biochemical characteristics [5]. These serovars are traditionally divided into typhoidal and non-typhoidal groups according to their host specificity and disease manifestations [6, 7]. Typhoidal *Salmonella* (TS), represented mainly by *S. enterica* serovars Typhi and Paratyphi A, B, and C, are highly specialized human-adapted pathogens responsible for enteric fever. Transmission occurs predominantly through the fecal-oral route via contaminated food or drinking water, although direct person-to-person transmission may also occur. After entering the host, these pathogens can evade early immune recognition and establish prolonged systemic infections characterized by persistent fever and bacteremia [5].

In contrast, non-typhoidal *Salmonella* (NTS) exhibit a broader host range and are commonly associated with zoonotic transmission. In most cases, NTS infections manifest as self-limiting gastroenteritis; however, some strains are capable of causing severe invasive disease [8]. These invasive non-typhoidal *Salmonella* (iNTS) strains penetrate the intestinal barrier and disseminate into the bloodstream or internal organs, resulting in septicemia and systemic infection. iNTS disease is particularly dangerous in infants, elderly individuals, malnourished patients, immunocompromised persons, and people living with HIV [9]. Mortality associated with bloodstream infections caused by iNTS remains high in many regions of the world [10]. Among the serovars frequently associated with invasive disease, *S. enterica* serovar Typhimurium is one of the most highly reported [11].

The introduction of molecular epidemiological approaches, particularly Multi-Locus Sequence Typing (MLST), has revealed extensive diversity within major *Salmonella* serovars [12]. MLST analysis demonstrated that strains belonging to the same serovar may form multiple clonal lineages differing in geographical distribution, host adaptation, virulence potential, and antimicrobial resistance profiles [13, 14]. Some sequence types (STs) are globally disseminated, whereas others appear to be restricted to specific ecological or geographical regions. For example, *S. Typhimurium* ST19 is considered one of the most widespread and evolutionarily generalized lineages, commonly associated with zoonotic transmission from livestock to humans [15]. In contrast, ST34 strains are frequently characterized by broader antimicrobial resistance profiles, including multidrug resistance, and have become increasingly associated with intensive animal production systems and foodborne transmission [16, 17].

Regional studies indicate that the population structure of *Salmonella* may differ substantially between countries and epidemiological settings [14]. In Kazakhstan, clinical isolates are predominantly represented by strains belonging to the ST11 genotype [18]. Nevertheless, the evolutionary relationships, virulence-associated genomic features, and epigenetic characteristics of these circulating strains remain insufficiently characterized.

Advances in third-generation sequencing technologies, including PacBio SMRT sequencing technology and Oxford Nanopore sequencing technology, now enable not only high-quality genome assembly but also direct detection of epigenetic modifications such as DNA methylation [19]. These approaches provide new opportunities to investigate the roles of restriction–modification systems, methyltransferases, horizontal gene transfer, and epigenetic regulation in the adaptation and pathogenicity of *Salmonella* strains [20, 21].

Nucleotide methylation occurs through the activity of methyltransferase (MTases), which are often components of restriction-modification (RM) systems that protect bacterial cells from bacteriophages and conjugative plasmids. However, orphan MTases are also found, and their role in the biology of pathogenic microorganisms remains incompletely understood. Scientific evidence indicates that genomic DNA methylation represents one of the mechanisms regulating gene expression and has a substantial influence on pathogenesis, the development of antibiotic resistance, and the ability of pathogens to survive in the environment [20–22].

The present study was aimed at sequencing *S. enterica* isolates collected in Kazakhstan from clinical patients and animals. The genomes of the selected strains were obtained through sequencing of genomic DNA using the Illumina sequencing platform and PacBio sequencing platform in order to perform genotyping and investigate the evolutionary dynamics of the *S. enterica* population in Kazakhstan.

## Materials and Methods

### Ethics approval

All procedures performed within this study complied with international and national ethical standards for biomedical research and were approved by the local Bioethics Committee of the M. Aikimbayev’s National Scientific Center for Especially Dangerous Infections (NSCEDI) (Protocol No. 2 dated 12 February 2024). Clinical material was obtained only after patients had signed written informed consent forms. The consent procedure was documented by the attending physicians of the participating medical institutions. All patient personal data were de-identified prior to laboratory analyses, and all investigations were conducted exclusively using anonymized samples. Animal biological samples were collected with the permission of the animal owners and in accordance with current biosafety and bioethics requirements.

### Culture isolation and growth conditions

The present study reports the results of a comprehensive characterization of 22 bacterial strains of the genus *Salmonella* isolated in Kazakhstan between 1957 and 2025. The study included isolates of diverse origins obtained from humans, animals, and collection repositories. Fourteen isolates were selected from the National Microorganism Collection (NMC) at NSCEDI (Almaty, Kazakhstan). Additionally, five clinical isolates obtained from the bacteriological laboratory of the I. Zhekenova City Clinical Infectious Diseases Hospital (CCIDH, Almaty, Kazakhstan) were included in the study. A further four isolates were provided by the Kazakh Scientific Research Veterinary Institute (KSRVI).

The investigated isolates cover a broad temporal range, diverse geographic regions, and multiple epidemiological sources of isolation. The dataset included both epidemiologically dominant serovars and rare genetic lineages of *S. enterica* circulating in Kazakhstan.

Strain cultivation was performed on Hottinger agar and meat-peptone agar (MPA) at 37°C for 24 h. The cultural and morphological characteristics of the isolates were evaluated according to standard microbiological protocols [26]. Growth dynamics and colony formation characteristics were monitored after 12, 24, and 48 hours of incubation using light microscopy and visual assessment of colony morphology. The analysis included determination of colony shape, transparency, surface characteristics, pigmentation, density, and diameter, as well as evaluation of the uniformity of bacterial growth.

The susceptibility/resistance of *S. enteridis* strains to antibiotics including amikacin (30 µg), ampicillin (10 µg), levofloxacin (5 µg), ceftriaxone (30 µg), and gentamicin (10 µg) was assessed in accordance with established methodological guidelines [23].

### Serological testing

Serological identification of the investigated isolates was performed using the agglutination reaction with polyvalent antisera of groups ABCDE, and monoreceptor Salmonella O- and H-antisera. Diagnostic antisera produced by the Saint Petersburg Research Institute of Vaccines and Sera and PETSAL (Russia) were used in this study. Serotyping was performed according to standard *Salmonella* serological classification principles based on somatic (O) and flagellar (H) antigen determination following the White-Kauffmann-Le Minor scheme [24].

The reference strain *S. enterica* pathovar Typhimurium TA98 (V RCM-0162) from the Russian Collection of Microorganisms (RCM) was used as a control microorganism in all experimental series and served for the standardization of bacteriological and molecular studies.

### DNA isolation and genome sequencing

Genomic DNA was extracted from pure cultures of *S. enterica* using the QIAamp DNA Mini Kit (Qiagen, Germany) according to the manufacturer’s protocol. To ensure high sample quality and remove potential inhibitors of enzymatic reactions, an additional purification step was performed using AMPure XP magnetic beads (Beckman Coulter, USA). The integrity and molecular weight of the extracted DNA were assessed by agarose gel electrophoresis. DNA concentration and purity were determined using a NanoDrop 1000 spectrophotometer (Thermo Fisher Scientific, USA) and a Qubit 4 fluorometer (Thermo Fisher Scientific, USA).

Libraries for whole-genome sequencing were prepared using the tagmentation-based Illumina DNA Prep kit (Illumina, USA) with unique dual indexes from the IDT for Illumina DNA/RNA UD Indexes Set A (IDT, USA). Sequencing was performed on an Illumina MiSeq platform (Illumina, USA) installed in NSCEDI using the MiSeq Reagent Kit v3 (600 cycles). A total of 17 pM DNA libraries were loaded into the sequencer for each run. Primary processing and quality control of the generated reads were carried out using the instrument’s standard software tools.

Five DNA samples were submitted to Inqaba Biotech (Pretoria, South Africa, https://inqababiotec.co.za/) for sequencing on the SMRT Pacific Bioscience (PacBio) Revio sequencing platform for epigenetic modification profiling.

### Bioinformatics tools and statistical analysis

De novo genome assembly was performed using the assemblers SPAdes v3.15.0 for short Illumina reads [25], and Flye v2.9.6 for long PacBio reads [26]. Plasmid clustering, reconstruction, and typing were carried out using MOB-suite v3.1.9 [27]. Integration of sequencing data and optimization of short-read assembly was performed using the ILLAMINA v1.0 pipeline (https://zenodo.org/records/15227656); and the LRASS v1.0 pipeline (https://zenodo.org/records/16571119) was used for the long-read assembly.

Identification of antimicrobial resistance genes was conducted using Abricate v1.4.0 (https://github.com/tseemann/abricate) and the CARD database through RGI v6.0.5 [28]. Heatmaps were generated in Python using json-2.0.9, panda-2.2.3, NumPy-2.3.2, Matplotlib-3.10.1, and SciPy-1.15.2 using gene presence-absence matrix generated from Abricate and RGI output files. Sample similarity was estimated using the Hamming distance applied to binary feature profiles, followed by hierarchical clustering using average linkage (UPGMA) implemented in scipy.cluster.hierarchy. Matrix rows were reordered according to dendrogram leaf order, and the resulting clustering tree (cladogram) was plotted adjacent to the heatmap.

Multilocus sequence typing (MLST) was performed based on the PubMLST online database (https://pubmlst.org/organisms/salmonella-spp) using the MLST scheme based on sequence types of seven core genes: *aroC*, *dnaN*, *hemD*, *hisD*, *purE*, *sucA*, and *thrA*. If the sequence type (ST) remained unidentified by the MLST scheme, the SalmcgMLST v1.0 core genome model was applied.

Programs ipdSummary and MotifMaker, parts of the software package smrtlink_25.2.0.266456 (https://www.pacb.com/support/software-downloads/?utm_source=chatgpt.com) installed on the computer cluster at the Centre for Bioinformatics and Computational Biology (University of Pretoria, Pretoria, South Africa), were used for statistical prediction of epigenetically modified nucleotides and DNA methylation motifs, respectively. For estimation of distribution of methylated sites across coding, non-coding and promoter regions (75 bp upstream of predicted gene transcription start codons), the assembled sequences were annotated using program Bakta v.1.9.4 [29].

Detection and analysis of restriction-modification (RM) gene clusters in assembled genomes was carried out using DefenseFinder and macsyfinder v2.1.4 [30]. Reconstruction of 3D structures of methyltransferase proteins by their amino acid sequences was conducted on ColabFold v1.6.1: AlphaFold2 Web-based notebook (https://colab.research.google.com/github/sokrypton/ColabFold/blob/main/batch/AlphaFold2_bat ch.ipynb) using MMseqs2 [31]. Program MEGA12 [32] was used for protein sequence alignment, identification of mutated loci and clustering proteins using the Neighbor-Joining algorithm. For visualization of predicted PDB protein structures and positioning of mutated contigs, Python 3 module py3Dmol-2.5.5 was used.

### Data availability

The sequenced genomes were deposited at the NCBI under BioProject PRJNA1468550.

## Results

### *S. enterica* strains used in this study

Collection *S. enterica* strains and recent hospital isolates were included in this study. The hospital strains were isolated from patients hospitalized with clinical manifestations of acute gastroenteritis. Four laboratory-confirmed cases of salmonellosis were identified. The patients ranged in age from 46 to 67 years and included three women and one man.

In all cases, the disease presented as a moderate gastrointestinal form of salmonellosis. The most common clinical manifestations included diarrhoea, abdominal pain, nausea, vomiting, fever and/or chills, general weakness, signs of dehydration, and loss of appetite. The frequency of diarrheal episodes ranged from more than five to approximately twenty episodes per day in the most severe cases. Physical examination revealed tenderness in the epigastric and periumbilical regions in most patients.

Analysis of the epidemiological history suggested a probable foodborne route of infection transmission. Reported possible exposure factors and susceptibility to antibiotics of the isolates are shown in Table 1.

**Table 1.** Epidemiological and microbiological characteristics of *S. enterica* strains isolated from patients hospitalized with salmonellosis in Almaty, Kazakhstan, 2025.

| Strain | Sex | Age | Epidemiological history | Antibiotic susceptibility* |  |  |  |  |
| --- | --- | --- | --- | --- | --- | --- | --- | --- |
|  |  |  |  | Amk | Amp | Lev | Cef | Gen |
| 16S | Female | 67 | Consumption of food during a festive event | S | R | S | R | S |
| 17S | Male | 49 | Chicken sandwich | S | R | S | S | S |
| 19S | Female | 46 | Raw egg | S | R | S | S | S |
| 20S | Female | 60 | Kazy (traditional horse meat sausage) | S | R | S | S | S |

\* Antibiotic abbreviations: Amk – amikacin; Amp – ampicillin; Lev – levofloxacin; Cef – ceftriaxone; Gen – gentamicin. S – susceptible; R – resistant.

None of the hospitalized patients reported recent travel outside the Republic of Kazakhstan, indicating a local origin of infection. Clinical presentations were consistent with the typical course of non-typhoidal salmonellosis with predominant involvement of the gastrointestinal tract. Although no severe complications or generalized forms of infection were observed, the severity of diarrheal symptoms and signs of intoxication required hospitalization and rehydration therapy.

### Phenotypic and bacteriological analysis of selected *S. enterica* strains

All 22 investigated *Salmonella* isolates, both collection and hospital strains, appeared as Gram-negative straight rods with rounded ends upon microscopic examination of Gram-stained smears. Cultivation on Hottinger agar and meat-peptone agar (MPA) resulted in the formation of smooth, transparent, round colonies with regular margins and a slight bluish tint. Colony diameters varied from 2 to 6 mm depending on incubation time and serovar characteristics. Growth was characterized by uniform colony morphology without signs of dissociation, indicating the stability of the cultural properties of the investigated strains.

The biochemical properties of the investigated isolates were determined using standard differential diagnostic media and test systems in accordance with the recommendations of Bergey’s Manual of Systematic Bacteriology. All investigated strains displayed a metabolic profile typical of members of the genus. The isolates fermented glucose, mannitol, maltose, dulcitol, sorbitol, arabinose, rhamnose, and xylose with the production of acid and gas, while lactose and sucrose were not fermented. All strains demonstrated growth on Simmons citrate medium, confirming their ability to utilize citrate as the sole carbon source. During cultivation, hydrogen sulphide (H_2_S) production and the ability to reduce nitrates to nitrites were observed. At the same time, all investigated strains lacked indole production and the ability to hydrolyse gelatine. The combination of biochemical characteristics confirmed the affiliation of the investigated cultures with the genus *Salmonella* and indicated the preservation of the characteristic metabolic phenotype among circulating strains.

All investigated isolates demonstrated positive agglutination reactions with polyvalent antisera of groups ABCDE, confirming their affiliation with the genus *Salmonella*. The results of identification of pathovars of *S. enterica* strains based on White-Kauffmann-Le Minor serotyping and MLST are shown in Table 2.

**Table 2.**
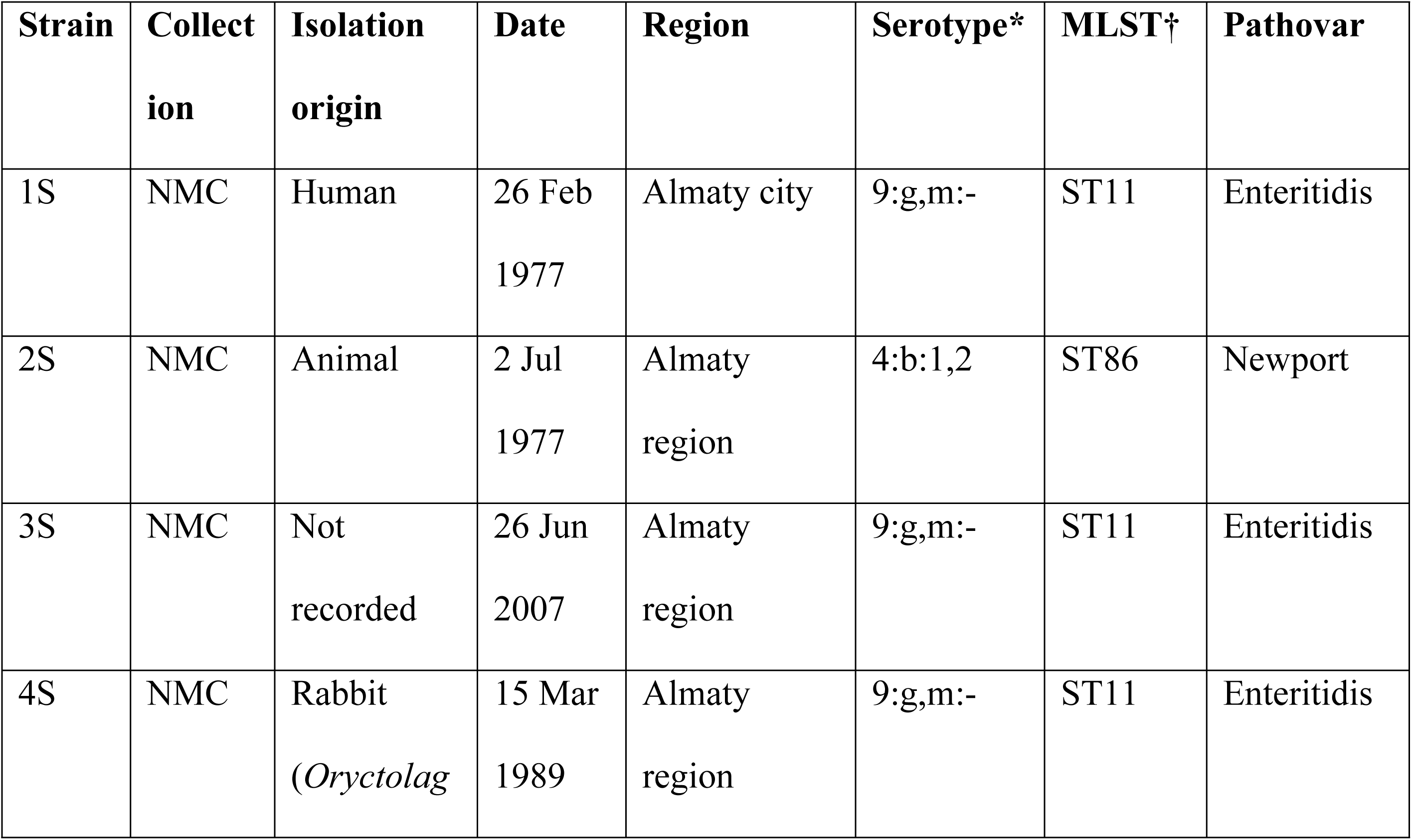
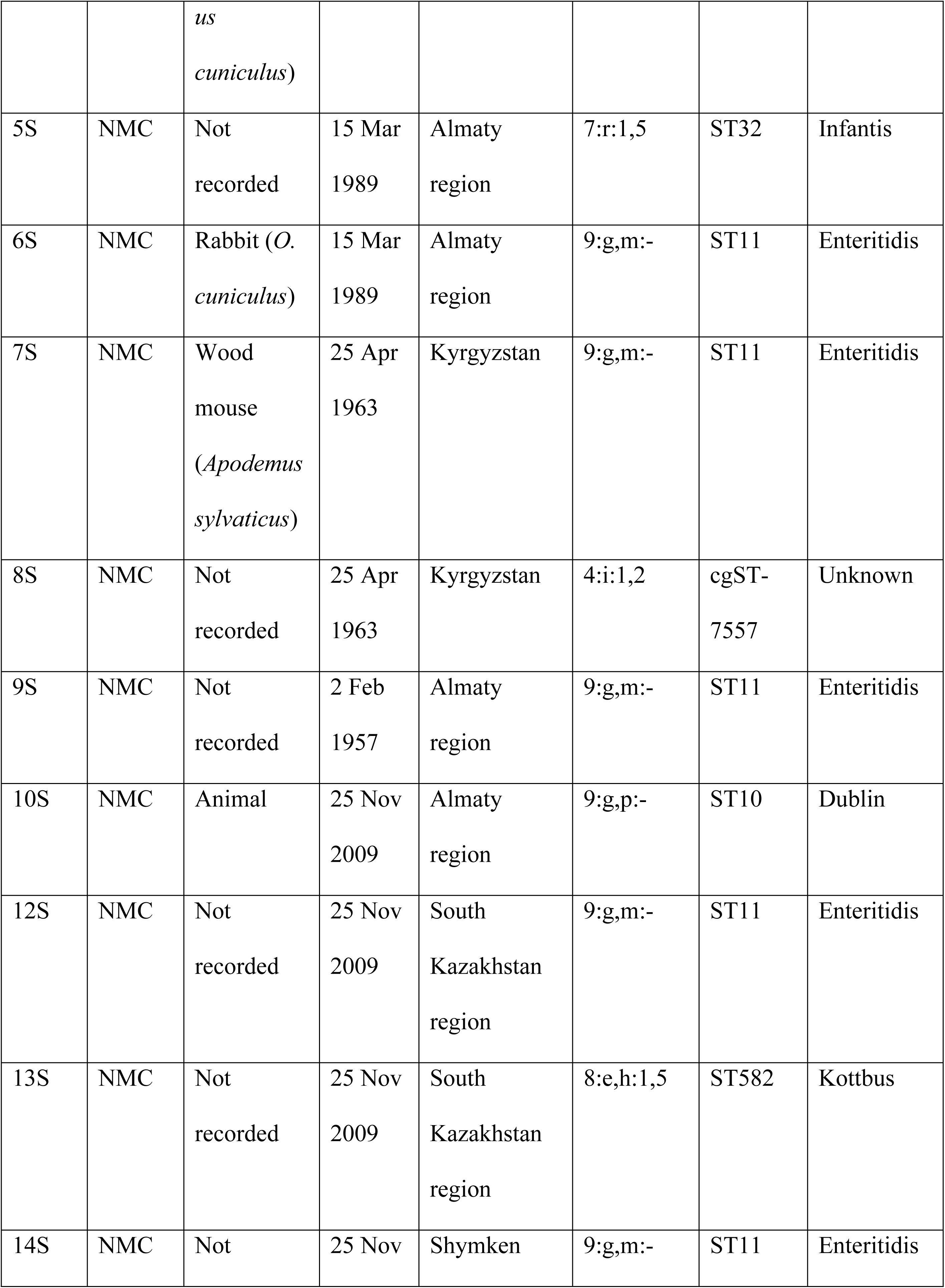
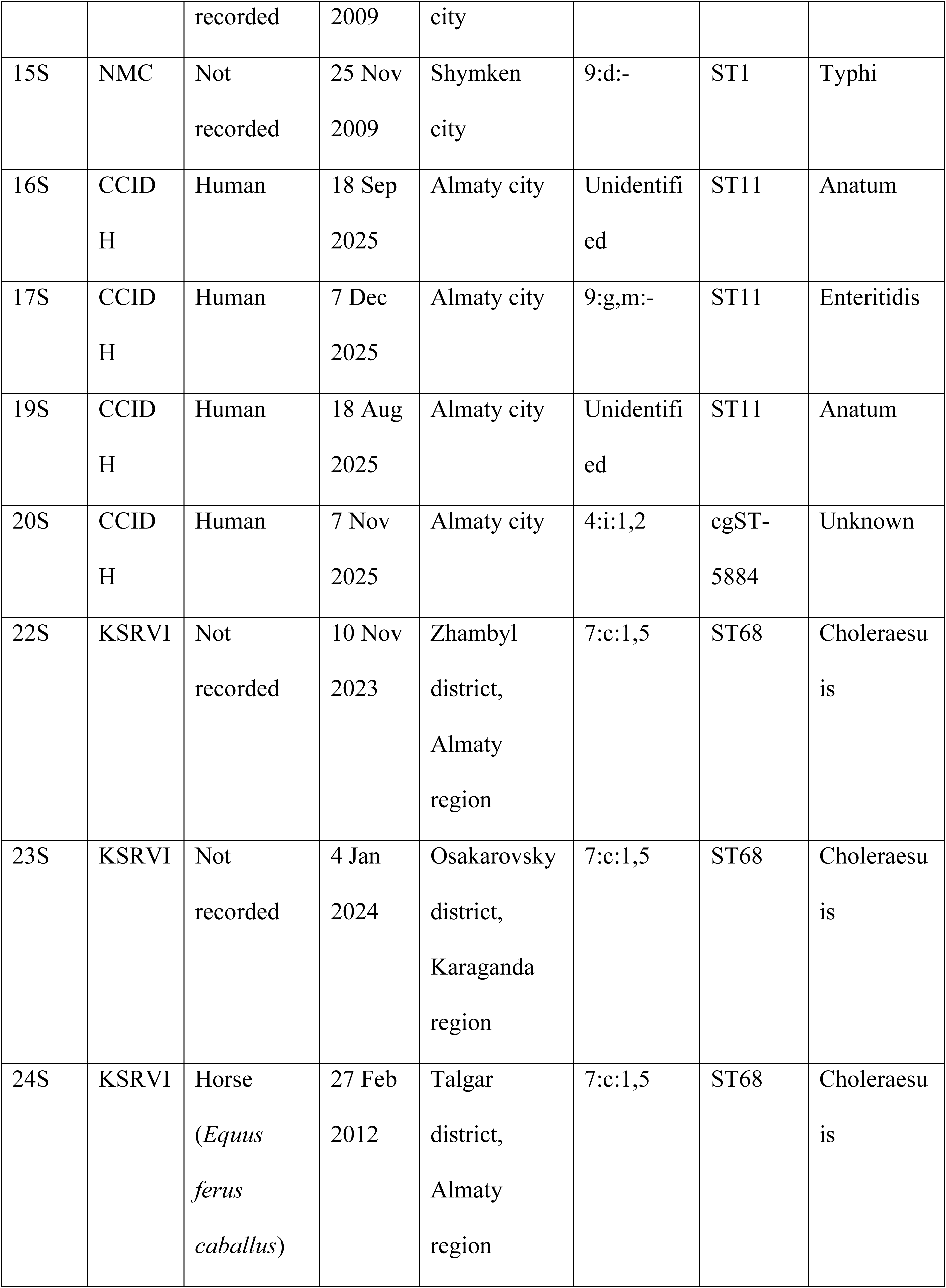
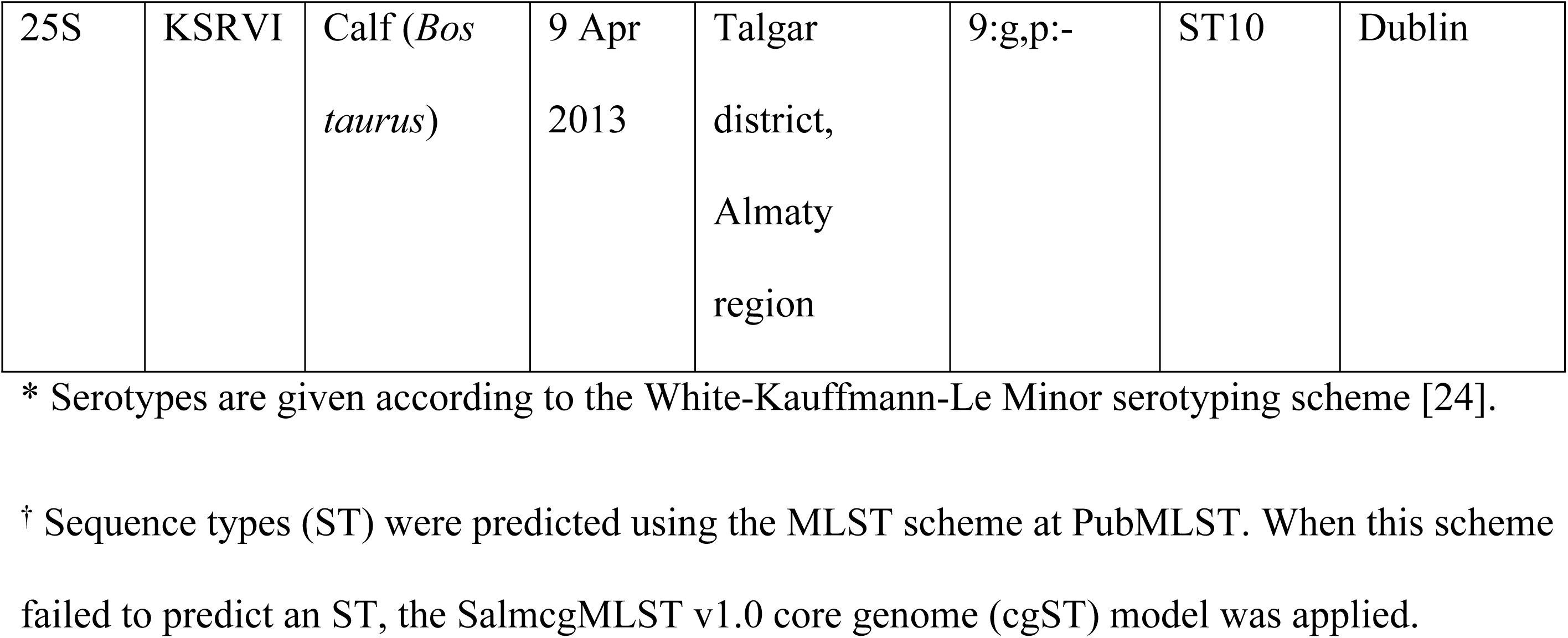
Properties and origin of assembled genome sequences of *S. enterica*.

### Genome sequencing

Whole-genome DNA sequences were obtained by sequencing the isolated samples on an Illumina sequencing platform. Five samples were additionally sequenced using the SMRT PacBio Revio sequencer to investigate the profiles of epigenetic modifications in genomic DNA.

To assemble DNA fragments generated by Illumina sequencing into contigs, the program SPAdes was used [25]. The MOB-suite software [27] was employed to identify plasmid contigs. For complete genome assembly, the ILLAMINA pipeline (https://zenodo.org/records/15227656) was applied. This pipeline integrates de novo genome assembly results with reference-based assembly approaches. The genome assembly results are presented in Table 3.

**Table 3.** Properties and origin of assembled genome sequences of *S. enterica*.

| Strain | Chromosome length | Chromosome GC-content | Gene number | Plasmid number | Plasmid gene numbers |
| --- | --- | --- | --- | --- | --- |
| 1S | 4884062 | 51.82% | 4550 | 3 | 93, 88, 3 |
| 2S | 4649739 | 52.27% | 4297 | 1 | 87 |
| 3S | 4884418 | 51.82% | 4557 | 3 | 92, 89, 1 |
| 4S | 4883554 | 51.82% | 4558 | 3 | 92, 88, 1 |
| 5S | 4881291 | 52.16% | 4508 | 1 | 87 |
| 6S | 4884114 | 51.82% | 4556 | 3 | 93, 88, 1 |
| 7S | 4883922 | 51.82% | 4551 | 2 | 93, 88 |
| 8S | 4964964 | 50.53% | 4698 | 1 | 85 |
| 9S | 4883893 | 51.82% | 4553 | 1 | 88 |
| 10S | 4886993 | 52.07% | 4624 | 3 | 91, 88, 48 |
| 12S | 4883793 | 51.82% | 4554 | 3 | 93, 88, 1 |
| 13S | 4884338 | 51.82% | 4537 | 4 | 87, 4, 3, 1 |
| 14S | 4884314 | 51.82% | 4553 | 3 | 92, 88, 3 |
| 15S | 4586611 | 52.21% | 4358 | 1 | 88 |
| 16S | 4882966 | 51.79% | 4554 | — | — |
| 17S | 4883897 | 51.82% | 4552 | 2 | 92, 88 |
| 19S | 4882774 | 51.75% | 4546 | 3 | 103, 49, 4 |
| 20S | 4964860 | 50.51% | 4693 | 1 | 95 |
| 22S | 4650091 | 52.24% | 4370 | 2 | 93, 87 |
| 23S | 4650096 | 52.24% | 4369 | 2 | 91, 87 |
| 24S | 4649113 | 52.24% | 4371 | 2 | 92, 87 |
| 25S | 4886157 | 51.71% | 4557 | 3 | 91, 87, 48 |

It was found that most of the selected *S. enterica* strains, with the exception of strain 16S, contained from one to five plasmids. A detailed characterization of the plasmid genomes is presented in Table 4.

**Table 4.** Plasmids identified in *S. enterica* strains, their genome lengths, and their assignment to different incompatibility groups.

| Strain | AF598 | AB232<br>(IncFII,<br>IncX1) | AD529 | AB241<br>(ColpVC) | AA974<br>(Col156) | AA474<br>(IncI1/B/<br>O) | AA619<br>(IncX4) |
| --- | --- | --- | --- | --- | --- | --- | --- |
| 1S | 70131 | 74672 | 3126 | — | — | — | — |
| 2S | 70173 | — | — | — | — | — | — |
| 3S | 70142 | 74654 | — | 2223 | — | — | — |
| 4S | 70113 | 74695 | — | 2228 | — | — | — |
| 5S | 70103 | — | — | — | — | — | — |
| 6S | 70160 | 74634 | — | 2257 | — | — | — |
| 7S | 70087 | 74677 | — | — | — | — | — |
| 8S | 70120 | — | — | — | — | — | — |
| 9S | 70114 | — | — | — | — | — | — |
| 10S | 70103 | 74655 | — | — | — | — | 32112 |
| 12S | 70174 | 74677 | — | 2240 | — | — | — |
| 13S | 70090 | — | 3144 | 2234 | 4648 | — | — |
| 14S | 70187 | 74622 | 3125 | — | — | — | — |
| 15S | 70115 | — | — | — | — | — | — |
| 16S | — | — | — | — | — | — | — |
| 17S | 70137 | 74674 | — | — | — | — | — |
| 19S | — | — | — | — | 4597 | 92347 | 32165 |
| 20S | — | 74705 | — | — | — | — | — |
| 22S | 70115 | 74724 | — | — | — | — | — |
| 23S | 70109 | 74724 | — | — | — | — | — |
| 24S | 70143 | 74746 | — | — | — | — | — |
| 25S | 70111 | 74684 | — | — | — | — | 32105 |

### Virulence and antibiotic resistance gene diversity

The search for virulence genes was performed against the VFDB database using the Abricate program. Antibiotic resistance genes were identified using the RGI tool. Presence-absence matrices of virulence and resistance genes across different strains are shown in Figures 1 and 2, respectively. Full lists of identified genes are given in respective supplementary Tables S1 and S2. Based on these matrices, cladograms representing similarity in gene patterns were constructed, and MLST sequence types identified for each strain are shown.

**Fig. 1.**
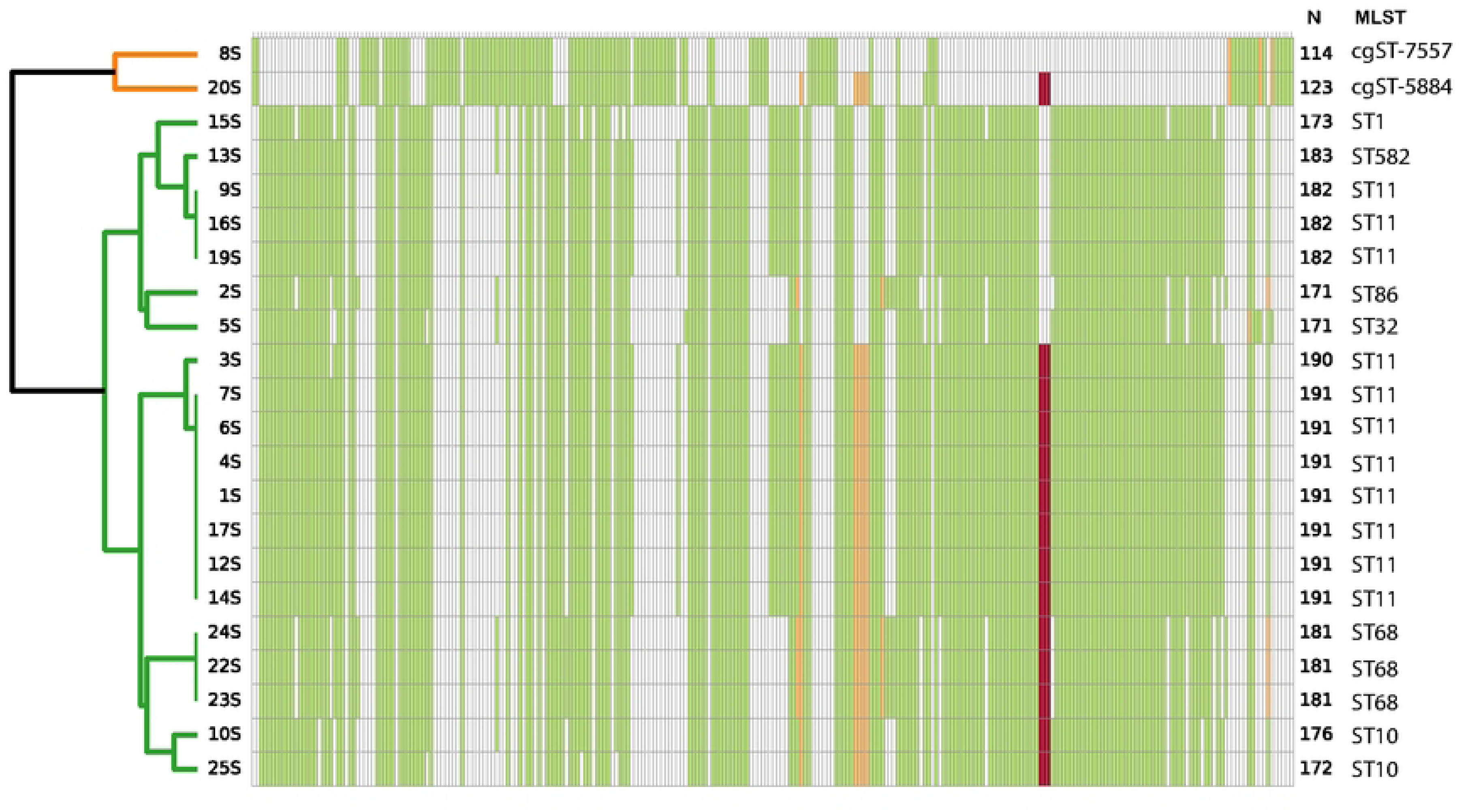
Presence-absence matrix of virulence genes in the genomes of sequenced *S. enterica* strains. White cells indicate absence of a given gene, while green cells indicate the presence of a single copy. Yellow and red cells denote the presence of two and three gene copies, respectively.

**Fig. 2.**
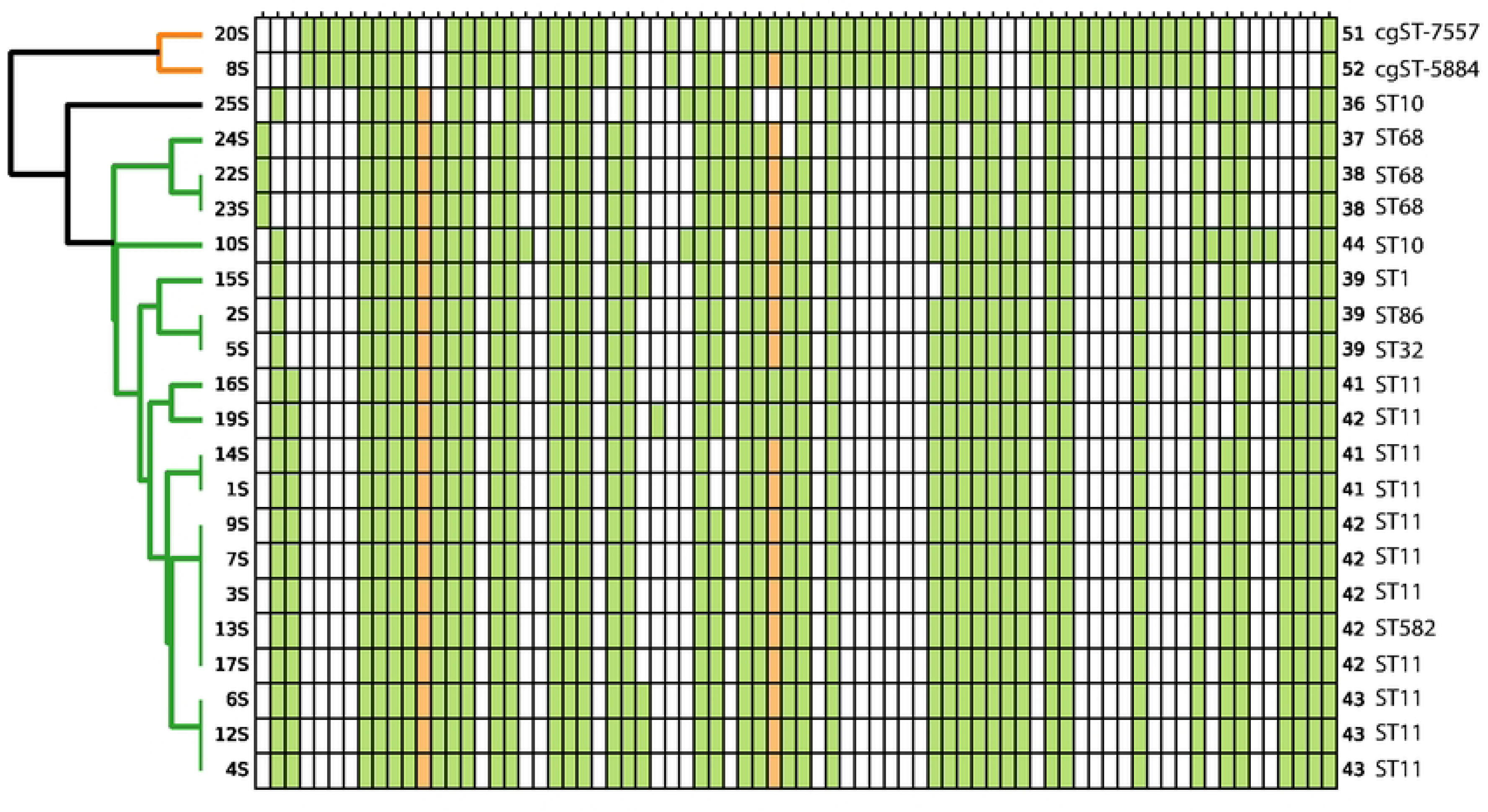
Presence-absence matrix of antibiotic resistance genes in the genomes of sequenced *S. enterica* strains. White cells indicate the absence of a given gene, while green cells indicate the presence of a single copy. Yellow cells denote the presence of two copies of the gene. Numbers on the right side of the plot indicate the total number of resistance genes identified in each genome. These numbers are followed by predicted MLST sequence types of the strains.

Numbers on the right side of the plot indicate the total number of virulence genes identified in each genome. These numbers are followed by predicted MLST sequence types of the strains.

A substantial proportion of virulence genes comprised those encoding secretion systems responsible for delivering toxins and effectors into host cells, as well as genes encoding the effectors themselves. In particular, the type II secretion system (T2SS) was represented by the genes *gspC–gspM*; the type III secretion system (T3SS) by the operons *invA–invJ, spaO–spaS, prgH–prgK,* and *orgA–orgC*, along with their transcriptional regulators *hilA, hilC, hilD,* and *invF*, and genes encoding toxic effectors such as *sipA, sipB, sipC, sipD, sicA, sicP, iagB, iacP, sopA, sopB (sigD), sopD, sopD2, sopE2, avrA, sptP,* and *slrP*.

The type IV secretion system (T4SS) was primarily plasmid-encoded, although in some strains it was also duplicated by chromosomal genes. Some strains of the ST11 clade additionally contained genes associated with the type VI secretion system (T6SS), including *acrE, acrF,* and *acrS*; however, several other genes required for a fully functional system were not detected.

Adhesin genes were also widely represented: *fimA–fimZ* (type 1 fimbriae); *lpfA–lpfE* (long polar fimbriae); *pefA–pefD* and *mrkA–mrkF* (other fimbrial systems commonly found in Enterobacteriaceae); *csgA–csgG* and *csgD* (curli biofilm proteins); as well as adhesins *bapA, fdeC, misL, shdA, pagN, rck, sinH, siiE,* and *yagV–yagZ*.

Iron acquisition and transport systems play a crucial role in microbial virulence, as iron ions are a limiting factor for bacterial growth during infection. The sequenced chromosomes contain genes involved in the synthesis of enterobactin (*entA–entF, entS*) and salmochelin (*iroB–iroN*), as well as genes encoding iron transport proteins (*fepA–fepG, fes*).

*Salmonella* strains produce and secrete systemic virulence factors such as *spvB, spvC,* and *spvD*, the cytolethal toxin *cdtB*, and the hemolysin *hly*. In typhoidal strains, the repertoire of virulence factors additionally includes the typhoid toxin operon containing the genes *pltA, pltB,* and *pltC*.

To counteract host defense mechanisms, pathogenic bacteria rely on oxidative stress response genes such as *sodCI*, and resistance to antimicrobial peptides produced by the host, mediated by the gene *ugd*. A number of genes are also involved in survival within macrophages, including magnesium ion transporters *mgtB* and *mgtC*, as well as macrophage survival genes *mig-14* and *mig-5*.

Most of the antibiotic resistance genes shown in Fig. 2 were located on chromosomes, with the exception of the *bla* TEM β-lactamase gene on plasmid AA474 in strain 19S. In contrast to plasmid-borne genes, the majority of chromosomal genes shown in Fig. 2 represent typical components of the resistome of Enterobacteriaceae and are not specifically associated with resistance to clinically used antibiotics, but rather include general resistance genes and regulatory elements controlling these genes.

From the perspective of acquired resistance to high doses of antibiotics, in addition to the plasmid-encoded TEM β-lactamase, important genes include *fosA7* (fosfomycin resistance), *sul1* (sulfonamide resistance), *tet(A)* and *tet(B)* (broad-spectrum resistance, including tetracyclines), and *aadA2, APH(3′)-Ia,* and *APH(3′)-IIa* (aminoglycoside resistance), although their presence in the genome alone does not necessarily indicate a broadly drug-resistant phenotype. More characteristic are mutations in target proteins that render them insensitive to antibiotics. In particular, well-known resistance-associated mutations were identified in the genes *gyrA* and *parC* (fluoroquinolone resistance), as well as *uhpT* (fosfomycin resistance).

### Patterns of nucleotide methylation and epigenetic modifications

To verify methylation patterns, six strains were selected for additional sequencing on the PacBio Revio sequencing platform.

The genomes of all strains were characterized by adenine methylation typical of Enterobacteriaceae on both DNA strands within the nucleotide sequence *GatC* (here and below, methylated nucleotides and the corresponding bases on the complementary strand are indicated in lowercase letters). This methylation is controlled by the *dam* methyltransferase (MTase) gene, which is not associated with restriction-modification (RM) systems. Two copies of this gene were identified on the chromosomes of the sequenced *S. enterica* strains. In addition, an extra copy of the *dam* methyltransferase gene was detected on plasmid AF598.

DAM methyltransferase is a dual-function enzyme that, in addition to adenine methylation, also catalyzes methylation of adjacent cytosine residues at motifs *cRGKGatC*, as well as at super-palindromic motifs *cRGKGatCMCYg* [33, 34]. Analysis of the *S. enterica* genomes confirmed cytosine methylation in the same sequences associated with *GatC* motifs. Whereas adenine methylation at *GatC* motifs approached 100%, the associated cytosine methylation occurred in only approximately 50% of *CRGKGatC* and *cRGKGatCMCYg* motifs across different genomic regions and strains, suggesting a possible role for this methylation in the regulation of gene expression [34]. Cytosine methylation patterns at *cRGKGATC* and *cRGKGatCMCYg* motifs for strain 1S are shown respectively in Fig. 3 A and B.

**Fig. 3.**
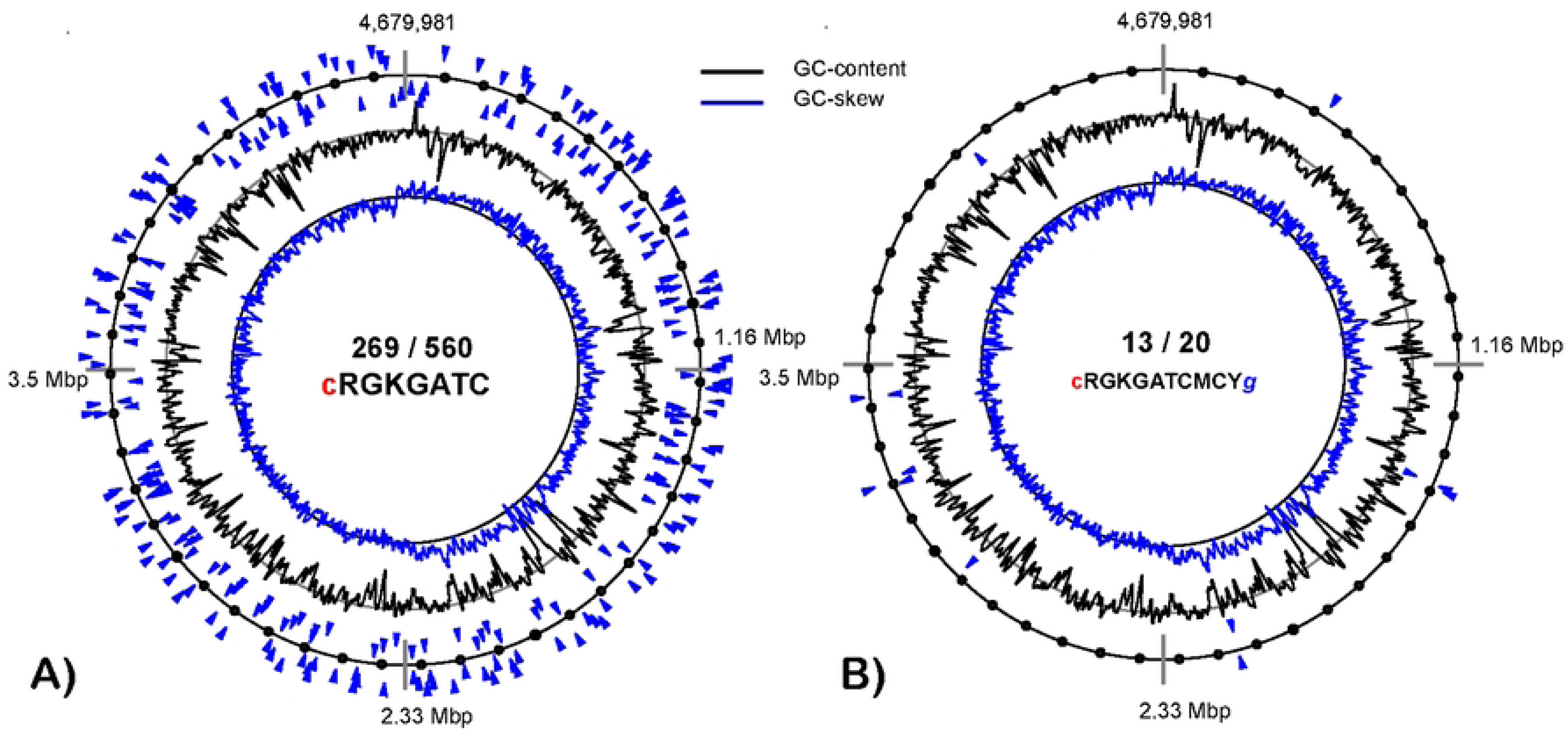
Patterns of cytosine methylation in strain 1S at A) *c*RGKGATC and B) *c*RGKGatCMCY*g* motifs.

DCM is another orphan MTase common in Enterobacteria that methylates cytosine residues in *CcWgG* motifs [21]. Genes encoding DCM cytosine-5-methyltransferase were found in all sequenced genomes, and genome 5S contains two copies of this gene, all located on the chromosomes. Although cytosine methylation at *CCWGG* motifs was abundant, with around 7,000 methylated sites identified in all genomes, this corresponded to only approximately 35% of the available methylation sites, leaving many cytosine residues within these motifs unmethylated.

An unexpected finding was the detection, in all genomes, of stable signals of epigenetic modifications involving cytosine in the semi-conservative motif *cCAGGBRD* and guanine in the motif *GNgGTVVCRD*. Although this cytosine methylation overlaps with the DCM-specific *CCWGG* motifs, the first cytosine in the motif is methylated, whereas DCM MTase methylates the second cytosine of this motif. The number of methylated *cCWGg* motifs was similar to the number of methylated *CcWgG* motifs, with around 7,000 sites per genome; however, these two methylation patterns did not overlap as no *ccWgg* methylated motifs were detected.

Epigenetic modifications of guanine are usually associated with oxidation to 8-oxoguanine; however, guanine oxidation typically occurs spontaneously and is not associated with specific sequence motifs. These guanine- and cytosine-associated epigenetic modifications were interrelated, since the motif *CCAGG* is complementary to the motif *GGTVV*. The *GNgGTVVCRD* sequences were found more frequently on the leading replichore of the chromosome, with a statistically significant correlation between motif frequency and GC-skew in sliding windows ranging from 0.10 to 0.17 (mean 0.13; p = 0.05). Modified guanine residues within these motifs were more frequent in coding regions (Z-score 2.36 ± 0.05, p = 0.018), while showing a random distribution across non-coding and promoter regions.

In contrast, the motif *cCAGGBRD* occurred more frequently on the lagging replichore, with correlation coefficients between the methylation frequency and GC-skew ranging from −0.16 to −0.08 (mean −0.12; p = 0.0). These motifs were more frequent in non-coding regions (Z-score 6.9 ± 0.03, p-value = 0.0) and less frequent in coding regions (Z-score −7.43 ± 0.04, p-value = 0.0).

While orphan DAM and DCM MTases are abundant, DNA methylation in bacteria is usually associated with the activity of MTases paired with restriction enzymes (REs), forming RM complexes. Methylation of genomic DNA at specific nucleotide sequences (motifs) protects native DNA from RE activity, whereas invading DNA from bacteriophages or conjugative plasmids remains unprotected and is degraded by REs. Thus, RM systems function as a form of “immunity” in bacterial cells against bacteriophages and conjugative plasmids [36].

The search for RM operons in the sequenced genomes was performed using the DefenseFinder program [30] (Table 5).

**Table 5.** Results of the identification of RM operons in *S. enterica* genomes using the DefenseFinder program.

| Strain | Type I | Type II | Type IIG_2 | Type III | Type IV |
| --- | --- | --- | --- | --- | --- |
| 1S | 1 | 0 | 0 | 1 | 0 |
| 2S | 1 | 0 | 0 | 1 | 0 |
| 3S | 1 | 0 | 0 | 1 | 0 |
| 4S | 1 | 0 | 0 | 1 | 0 |
| 5S | 1 | 1 | 0 | 1 | 0 |
| 6S | 1 | 0 | 0 | 1 | 0 |
| 7S | 1 | 0 | 0 | 1 | 0 |
| 8S | 1 | 0 | 0 | 0 | 2 |
| 9S | 1 | 0 | 0 | 1 | 0 |
| 10S | 1 | 2 | 1 | 1 | 0 |
| 12S | 1 | 0 | 0 | 1 | 0 |
| 13S | 1 | 0 | 0 | 1 | 0 |
| 14S | 1 | 0 | 0 | 1 | 0 |
| 15S | 0 | 1 | 0 | 1 | 0 |
| 16S | 1 | 0 | 0 | 1 | 0 |
| 17S | 1 | 0 | 0 | 1 | 0 |
| 19S | 1 | 0 | 0 | 1 | 0 |
| 20S | 1 | 0 | 0 | 0 | 2 |
| 22S | 1 | 0 | 0 | 1 | 0 |
| 23S | 1 | 0 | 0 | 1 | 0 |
| 24S | 1 | 0 | 0 | 1 | 0 |
| 25S | 1 | 2 | 1 | 1 | 0 |

Two RM operons belonging to types I and III were identified in most genomes. Type I RM systems were detected in all strains. Type III RM systems were absent in strains 8S and 20S. Instead, these two strains each contained two alternative type IV RM operons. The largest diversity of RM operons was observed in genome 25S, which contained RM systems of types I, II (two operons), IIG-2, and III. It should be noted that the presence of RM operons on the chromosome does not guarantee their activity in genome methylation or phage defense. The methylation patterns were verified in six strains sequenced on the PacBio Revio sequencing platform.

Intensive adenine methylation at *CAGaG* motifs was detected in all sequenced genomes and most likely is associated with the type III RM system, the genes of which were also identified in all six sequenced strains. The methylated motifs were predominantly located in non-coding regions of the genome (Z-score 8.98 ± 0.02, p = 0.0) and in promoter regions of genes (Z-score 2.31 ± 0.08, p = 0.021), but were rarely found within coding regions of the genome (Z-score −10.59 ± 0.02, p = 0.001).

Genes encoding type I RM systems were identified in all sequenced strains (Table 5). However, analysis of epigenetic modification patterns revealed type I specific adenine methylation on both DNA strands at the motif *GGYaNNNNNNtCG* only in strain 19S (Fig. 4A). In the other five strains, no more than 5% of *GGYANNNNNNTCG* motifs were methylated, as illustrated for strain 1S in Fig. 4B, which can be considered as a residual or background methylation. In strain 19S, this methylation predominantly targeted coding sequences (Z-score 3.24 ± 0.03, p = 0.001) and was rare in promoter regions (Z-score −6.36 ± 0.1, p = 0.0), while showing a random distribution in non-coding regions.

**Fig. 4.**
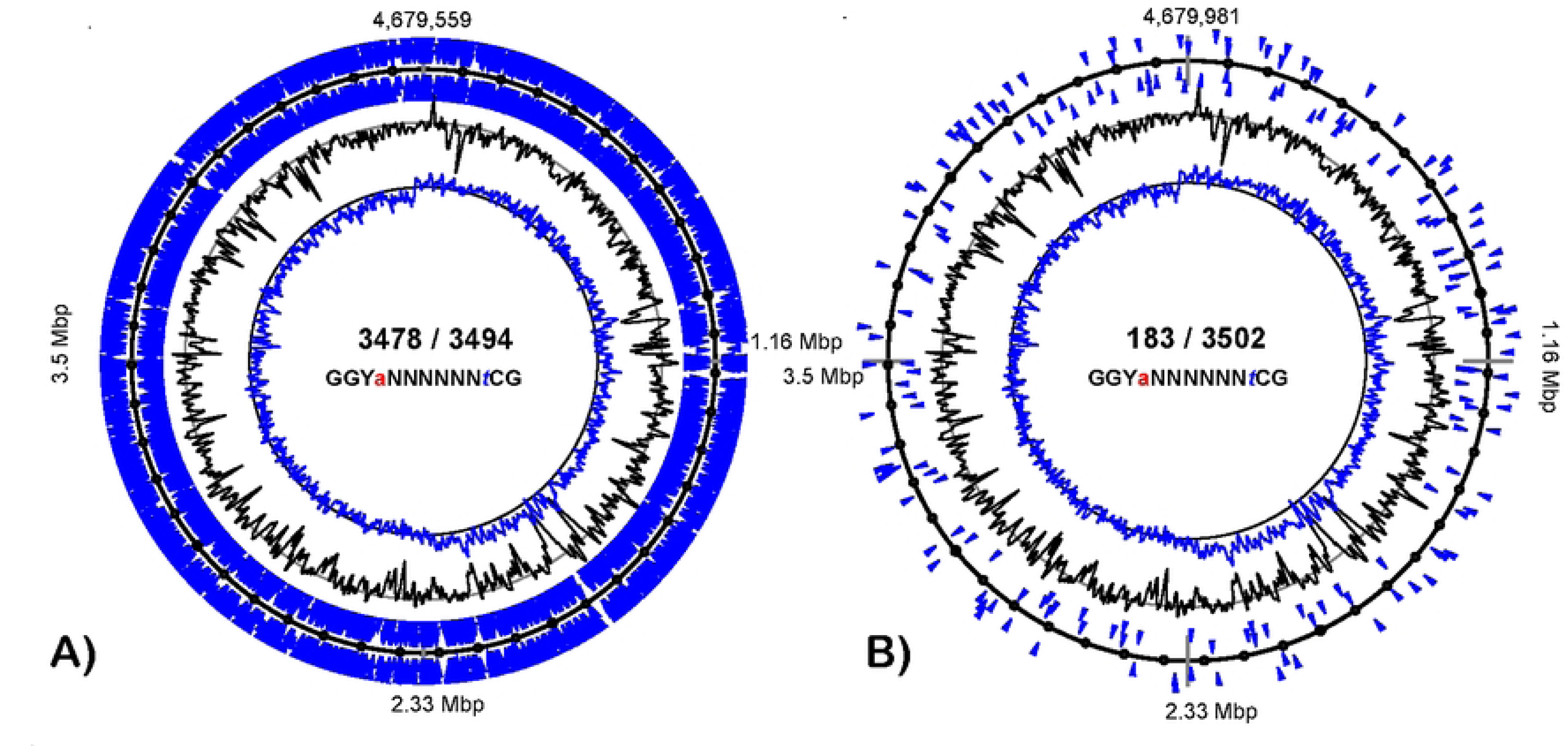
Patterns of adenine methylation at GGYaNNNNNNtCG motifs in A) 19S and B) 1S strains.

## Discussion

Analysis of the genomes of the selected strains confirmed that *S. enterica* isolates of the ST11 genotype are the most prevalent in clinical settings in Kazakhstan [18].

A large number of plasmids shown in Tables 3 and 4 indicates their significant role in the biology and survival strategy of *S. enterica*. Most of the isolated cultures contained two plasmids: AF598 and AB232. Plasmid AF598 carries several genes and operons important for bacterial metabolism. For instance, the operon comprising the genes *rhaT, rhaR, rhaS, rhaA, rhaB, rhaD,* and *rhaM* provides a complete pathway for L-rhamnose utilization. L-rhamnose metabolism is common among bacteria inhabiting plant-associated environments, where rhamnose-containing polysaccharides are abundant [37]. This suggests that this plasmid may play an important role in the long-term survival of the pathogen in the external environment, particularly in rhizosphere. Stress response genes present on this plasmid, such as *cpxA, cpxR, cpxP,* and *sodA*, are typically not directly associated with virulence, but may also contribute to environmental persistence.

In contrast, another large plasmid, AB232, present in most of the sequenced strains, is clearly associated with pathogen virulence. This plasmid carries a complete set of genes encoding a type IV secretion system (T4SS), including *virD4, virB4, virB9, virB11,* and *ptlE*. In addition, the genes *spvB* and *spvC* encode virulence factors responsible for the destruction of host cell cytoskeletal microtubules and suppression of the immune response, respectively. Other virulence-associated factors detected in this plasmid include the genes *rfaH, vsdE, mkaB,* and *mkaC*, as well as an invasin gene that mediates adhesion of the pathogen to host cells.

Furthermore, the plasmid contains conjugation genes and numerous transposons and integrases, facilitating plasmid transfer between cells and the integration of genetic material into the core genome of the pathogen. The absence of plasmid AB232 may indicate reduced virulence; however, in strain 19S this plasmid was replaced by a smaller plasmid carrying a similar repertoire of virulence genes, thereby maintaining pathogenicity of the strain while potentially increasing growth rate.

Other plasmids were specific to individual strains. For example, strain 19S, which lacked the two aforementioned plasmids, harbors a large plasmid AA474, in which the *bla* gene encoding TEM β-lactamase was identified. The *cia* colicin gene found in this plasmid is important for the displacement of competing intestinal microbiota during infection. Other genes on this plasmid are involved in conjugation and horizontal gene transfer. The same strain also carries another medium-sized conjugative plasmid, AA619, which is also present in strains 10S and 25S. This plasmid contains the genes *virB4, virB8, virB9,* and *virB11* of the type IV secretion system (T4SS), which is involved in pathogenesis. In strains 10S and 25S, these genes duplicate the T4SS genes found on the large plasmid AB232.

Strain 19S also contains a small plasmid, AA974, which is also found in strain 13S. This plasmid carries four genes of unknown function. Two other small plasmids, AD529 and AB241 (Table 4), likewise do not contain functional genes except for the regulatory gene *rop*, which is involved in the replication of ColE1-type plasmids.

Clustering of strains based on virulence and antibiotic resistance gene patterns shown in Figures 1 and 2, demonstrated that strains belonging to the same genotype (MLST ST) and serotype (Table 2) are not always homogeneous with respect to the presence or absence of genetic factors associated with horizontal gene transfer. Distinct combinations of virulence and resistance genes arise within bacterial clades as a result of horizontal gene transfer, natural selection in specific ecological niches, as well as differences in microbial survival strategies and predominant routes of transmission. In general, representatives of the less common genotypes (ST1, ST82, ST86, ST32, ST68, and ST10) were characterized by a reduced repertoire of virulence factors, which likely resulted in lower invasiveness.

Strains 8S and 20S were characterized by a conserved repertoire of virulence factors. These same strains also exhibited a unique set of resistance genes (Fig. 2), with a predominance of genetic factors involved in the regulation of gene expression, which may indicate adaptation to a more dynamically changing environment. Differences in resistance gene patterns among the strains mainly involved broad-spectrum efflux pumps, as well as the *sul1* and *tet(A/B)* genes, suggesting frequent horizontal transfer of these determinants.

In recent years, an increasing number of studies have suggested that changes in genome methylation patterns may directly or indirectly influence pathogen virulence and antimicrobial resistance [35, 38, 39]. MTase genes and entire RM operons are transferred between bacteria through horizontal gene exchange mediated by plasmids or bacteriophages [40]. Such exchange not only provides bacteria with new defense mechanisms against phages, but also affects patterns of gene expression and regulation through the emergence of new methylated nucleotide sites within genes and promoter regions [41].

Acquired RM operons gradually degenerate as a result of mutations. Consequently, bacterial genomes often contain more individual MTases and RM operons than detectable DNA methylation motifs [33]. The degradation of RM systems usually begins with an inactivation of RE, causing MTases to lose tight control over their activity and specificity. Degradation of RM systems makes bacterial strains more vulnerable to phage attack, but at the same time increases the likelihood of gene exchange via plasmids and transposons [33]. Invasive strains are relatively well protected from phages due to their intracellular parasitic lifestyle, whereas the acquisition and exchange of virulence factors and antibiotic resistance genes become critically important for their evolutionary success.

Because *Salmonella* is characterized by a dual lifestyle involving host invasion and hospital-associated infections, as well as persistence in water and plant rhizospheres populated by highly competitive microorganisms and bacteriophages [42–44], these bacteria require different survival strategies across highly diverse ecosystems. This may explain the presence of the two plasmids, AB232 and AF598, in the majority of the sequenced strains, as they provide genes required either for virulence or for environmental survival. It can be hypothesized that RM systems may also be fine-tuned for adaptation to different environments, while RM-associated and orphan MTases may participate in the regulation of gene expression [21].

The unexpected lack of DNA methylation activity at type I motifs in the sequenced genomes, despite the abundance and high level of conservation of the type I RM operon, may represent an example of such tightly regulated RM systems, which so far have been poorly characterized in the literature.

The type I RM gene cluster in the majority of sequenced *S. enterica* strains comprises four genes: the *mrr* RM-associated gene located on the opposite strand relative to the other genes of the RM operon; the *hsdR* type I RM endonuclease; the *hsdM* N6-DNA MTase; and the *hsdS* motif-recognition specificity subunit. Gene *mrr* was not present in type I RM operon of 8S and 20S genomes, but their operons contain two additional genes: ATP-dependent RNA helicase *srmB* and *dprA* gene encoding a DNA processing protein. There were several independent evolutionary origins of this operon found in different strains (Fig. 5 A-D) with HsdS motif-recognition subunit being the most variable between groups that makes it an optimal marker gene to distinguish between *S. enterica* lineages. However, within ST groups, the operons demonstrated strong conservation with no nucleotide substitutions detected in either coding or intergenic regions.

**Fig. 5.**
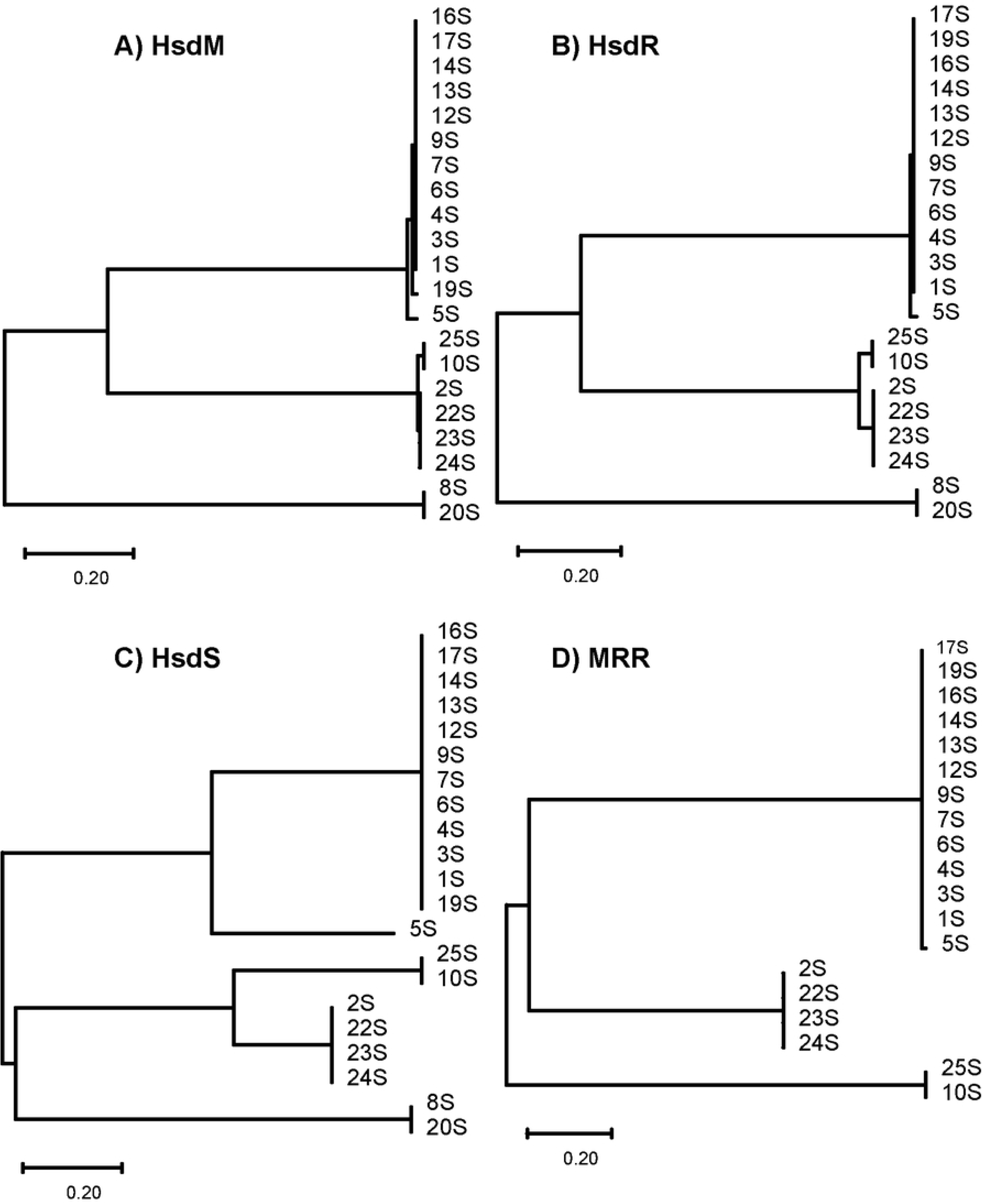
Neighbour-joining cladograms showing the sequence similarity of proteins encoded by type I RM operons: A) HsdM; B) HsdR; C) HsdS; and D) MRR.

An exception was observed in strain 19S, which differs from the other strains of the ST11 genotype by multiple nucleotide substitutions in *hsdM*, resulting in seven amino acid substitutions within two variable regions: (L103I, E105A, Y109N, A112S, and N119A) and (T363A and H366N). In this notation, the first letter represents the amino acid state in the majority of ST11 strains, the last letter represents the amino acid state in strain 19S, and the number indicates the codon position.

Surprisingly, mutant strain 19S was the only strain exhibiting detectable adenine methylation activity at type I motifs. Projection of the mutated codons onto the predicted three-dimensional structure of the HsdM protein (Fig. 6) showed that these substitutions are located opposite the DNA-binding helix–turn–helix motif. Further investigation is required to determine the possible role of these domains in the regulation of MTase activity, which could explain the active chromosomal methylation observed in the mutant strain as a result of escape from MTase suppression by a repressor.

**Fig. 6.**
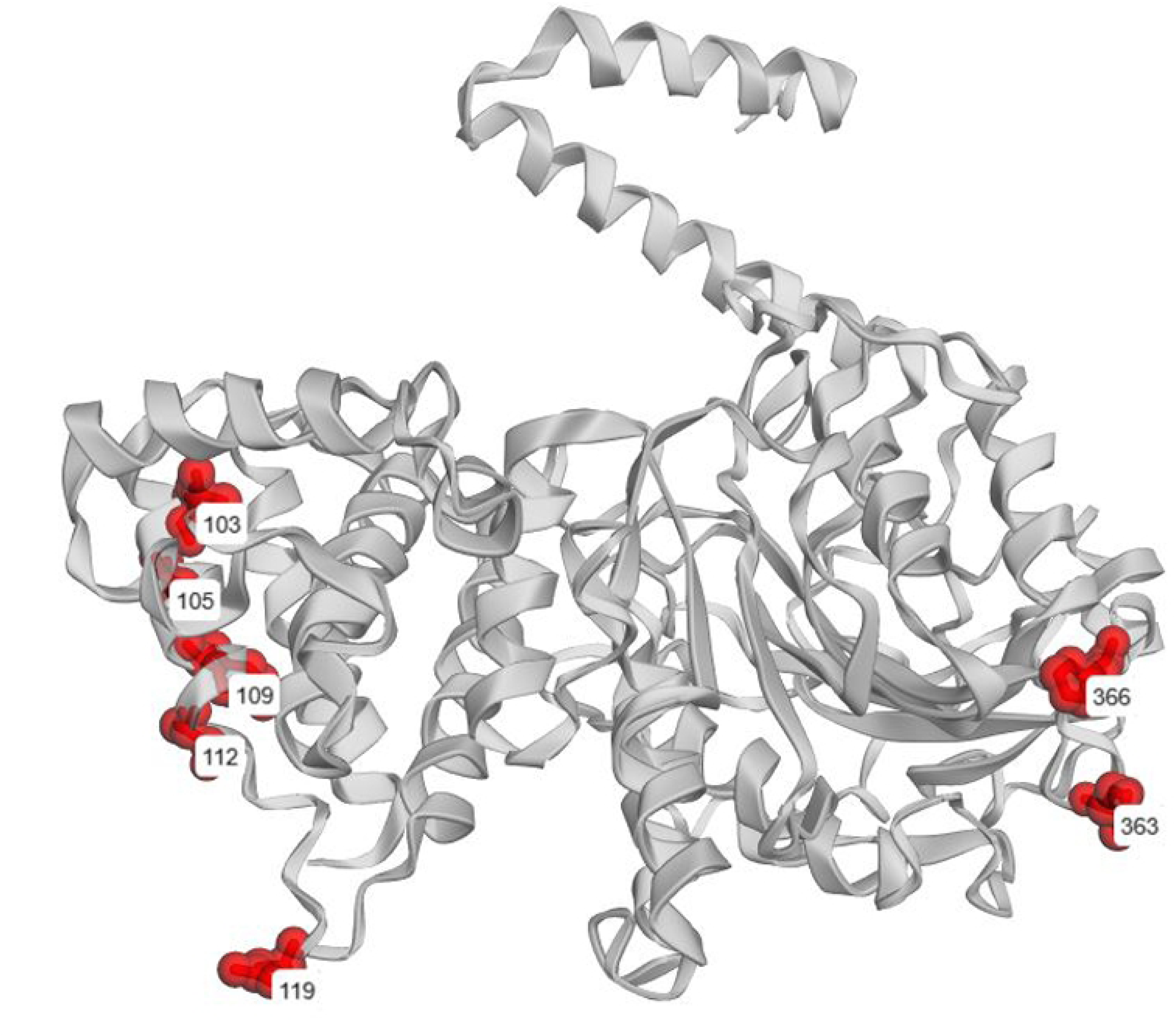
Projection of the mutated codons onto the predicted three-dimensional structure of the HsdM protein.

Activation in MTase activity, in turn, leads to a redistribution of methylated DNA regions. For example, in the genome of strain 19S, which possesses a super-active HsdM MTase, important genes such as the RNA polymerase gene *rpoB*, the methionine synthase gene *metH*, and the transmembrane transporter gene *xylP* became hypermethylated, each containing five methylation sites. In the other genomes, these genes remained unmethylated. Likewise, the promoter regions upstream of the *clpX* protease gene and the glutathione permease gene *gsiC* also remained unmethylated in those strains, but methylated in 19S.

Hospital strain 19S, isolated in 2025, may represent the emergence of a new lineage derived from *S. enterica* ST11 pathogens commonly found in Kazakhstan. This strain has completely reshaped its plasmid repertoire (Table 4) by replacing the large AF598 and AB232 plasmids, which are common to all other ST11 strains, with shorter plasmids enriched in virulence determinants and the *bla* TEM β-lactamase. Mutations in HsdM that activated MTase activity were most likely evolutionarily selected. Notably, all other genes, including the intergenic spacer regions within the type I RM gene cluster (*mrr* | *hsdR–hsdM–hsdS*) remained intact (Fig. 5). This observation suggests that restriction activity mediated by HsdR likely remained suppressed.

Therefore, the likely consequence of MTase activation was not enhanced protection against phages through the RM system, but rather regulatory genomic DNA methylation with the potential to influence gene expression patterns. This hypothesis should be validated in future transcriptomic studies.

DAM-mediated methylation at *GATC* motifs was more stable and approached 98% of all detected motifs. In the current study, notable differences were observed in the GATC methylation of promoter regions upstream of the *prfB-2* and *yqcC* genes, where methylation was absent in strains 2S and 10S, but three methylation sites were present in the other four strains. Previous studies have indicated that methylation within promoter regions can influence gene expression [33, 38, 40].

In contrast to the DAM methylation at adenine residues, DCM-mediated cytosine methylation at *CcWgG* motifs was partial, reaching approximately 35% of the available motifs. Variations in non-RM methylation of *S. enetrica* serovar Typhimurium genomes during different growth stages was reported [21]. Another study showed that DCM-mediated methylation at CCWGG motifs in *Yersinia pestis* is temperature-dependent and may be involved in gene regulation during the transition from vector insects to warm-blooded hosts [45]. A similar number of methylated *cCWGg* sites was detected in each genome; however, these two methylation patterns did not overlap. Significant differences in frequencies of *CcWgG* and *cCWGg* methylation, ranging from no methylation to 10-13 methylated sites per gene body, were observed across the sequenced genomes in the *rpoC* DNA-directed RNA polymerase beta-subunit, *acrB*_2 multidrug efflux pump, and several conserved hypothetical genes. Differential methylation at these motifs was also observed in the promoter regions of the following genes: *pdeN*_2; cyclic di-GMP phosphodiesterase, *lolA* outer-membrane lipoprotein carrier protein, *lldD* L-lactate dehydrogenase, *yecS*_3 L-cystine permease, *yejB* inner membrane ABC permease, and in *dcuR*/*dcuS* transcriptional regulators.

It is plausible that the same DCM MTase may be responsible for both methylation patterns, with its substrate specificity influenced by the surrounding nucleotide context. For example, *cCAGGBRD* methylation was most frequently observed when the *CCAGG* motif was followed by a *BRD* sequence. However, the detection of epigenetic modification signals involving cytosine and guanine within the tandem repeats *cCAGGBRD* and *GNgGTVVCRD* suggests that epigenetic modifications associated with CCWGG motifs may arise from an overlap of several independent processes.

Tandem repeats of semi-conservative motifs unevenly distributed between bacterial chromosome replichores have previously been described as architecture-imparting sequences (AIMS), which serve as binding sites for proteins involved in DNA replication and repair processes, such as FtsK [46]. Earlier studies have shown that AIMS repeats can be methylated in some bacteria [34]. It can therefore be hypothesized that interactions with replication and repair proteins in *S.enterica* may induce structural changes in DNA that are recognized by the PacBio sequencing platform as epigenetic modifications. The functional impact of these changes on genome activity requires further investigation.

## Conclusions

*S. enterica* remains a frequent causative agent of acute intestinal infections, often with a tendency toward systemic multiple organ infection [1, 8–11]. It is important to be able to identify sources of disease outbreaks in a timely manner and to accurately assess the epidemiological risks associated with disease spread. MLST genotyping enables the identification of genetic variants (clades and lineages) of pathogens belonging to the same species. However, it should be taken into account that MLST genotyping is based on the analysis of neutral mutations in core genes. These mutations usually do not carry direct biological significance and primarily serve as markers of common ancestry. Additional information on strain virulence and pathogenicity can be obtained by analyzing combinations of accessory genes, including virulence factors and antibiotic resistance genes.

In the present study, important differences were identified in the repertoires of virulence and antimicrobial resistance genes among *S. enterica* strains isolated in Kazakhstan from clinical samples, animals, and environmental sources. The ST11 genotype predominated among the isolates; however, strains of this genotype were separated into several genetically distinct clusters, likely reflecting their association with different epizootic outbreaks or divergent evolutionary trajectories of the pathogen.

The presence of two large plasmids, AF598 and AB232, in most of the studied strains suggests adaptation to invasive virulence combined with the ability to persist for extended periods in the environment, whereas the absence of one of these plasmids may reflect the predominance of either a parasitic or a saprophytic survival strategy. Of particular concern is the detection of a newly emerged sub-lineage derived from the ST11 lineage, represented by the recent hospital isolate 19S, which carries novel plasmids enriched in virulence and antibiotic resistance determinants.

Additional insights into the genetic variability of bacterial cultures can be obtained using modern sequencing technologies such as PacBio SMRT sequencing technology, which allows profiling of epigenetic modifications in microbial genomes.

The detection of MTase genes on bacterial chromosomes does not guarantee their functional activity, whereas DNA methylation profiles provide clear evidence regarding the number of active and degrading or suppressed RM systems. The presence of multiple active RM systems protecting bacterial cells against numerous bacteriophages and conjugative plasmids is usually associated with adaptation to environmental persistence, whereas suppression of these systems, which increases the frequency of horizontal genetic exchange, is more characteristic of invasive strains. Furthermore, differences in the repertoire and activity of RM systems and individual MTases enable more precise discrimination between bacterial clades within the same genotype through differences in methylation profiles. The influence of methylation and other types of nucleotide epigenetic modifications on gene expression in *S. enterica* remains an important subject for future investigation using transcriptomic approaches. The *S. enterica* strains sequenced in this project may serve as optimal model organisms for future studies on the role of DNA methylation and other epigenetic nucleotide modifications in the regulation of gene expression in pathogens, particularly strain 19S, which represents a natural mutant exhibiting increased DNA methylation at type I motifs affecting numerous genes and promoter regions.

---

**S1 Table**. Matrix of presence (1) and absence (0) of virulence genes in sequenced genomes.

**S2 Table.** Matrix of presence (1) and absence (0) of antibiotic resistance genes and mutations in sequenced genomes.

